# Pemivibart is less active against recent SARS-CoV-2 JN.1 sublineages

**DOI:** 10.1101/2024.08.12.607496

**Authors:** Qian Wang, Yicheng Guo, Jerren Ho, David D. Ho

## Abstract

Protection from COVID-19 vaccination is suboptimal in many immunocompromised individuals. In March 2024, the Food and Drug Administration issued an Emergency Use Authorization for pemivibart (Permagard/VYD222), an engineered human monoclonal antibody, for pre-exposure prophylaxis in this vulnerable population. However, SARS-CoV-2 has since evolved extensively, resulting in multiple Omicron JN.1 sublineages. We therefore evaluated the in vitro neutralizing activity of pemivibart against the prevalent forms of JN.1, including KP.2, KP.3, KP.2.3, LB.1, and, importantly, KP.3.1.1, which is now expanding most rapidly. A panel of VSV-based pseudoviruses representing major JN.1 sublineages was generated to assess their susceptibility to pemivibart neutralization in vitro. Structural analyses were then conducted to understand the impact of specific spike mutations on the virus-neutralization results. Pemivibart neutralized both JN.1 and KP.2 in vitro with comparable activity, whereas its potency was decreased slightly against LB.1, KP.2.3, and KP.3 but substantially against KP.3.1.1. Critically, the 50% inhibitory concentration of pemivibart against KP.3.1.1 was ∼6 µg/mL, or ∼32.7 fold higher than that of JN.1 in our study. Structural analyses suggest that Q493E and the S31-deletion mutations in viral spike contribute to the antibody evasion, with the latter having a more pronounced effect. Our findings show that pemivibart has lost substantial neutralizing activity in vitro against KP.3.1.1, the most rapidly expanding lineage of SARS-CoV-2 today. Close monitoring of its clinical efficacy is therefore warranted. These results also highlight the imperative to expand our arsenal of preventive agents to protect millions of immunocompromised individuals who could not respond robustly to COVID-19 vaccines.

## Main text

Pemivibart (Permagard; VYD222) was authorized in March 2024 for emergency use as pre-exposure prophylaxis against COVID-19 for immunocompromised individuals who could not respond robustly to vaccines^1^. This human monoclonal antibody was derived from ADG-2, an antibody directed to the SARS-CoV-2 spike [receptor-binding domain (RBD) class 1/4 region] that has previously demonstrated protective efficacy against SARS-CoV-2 infection^2^. However, ADG-2 lost virus-neutralizing activity against the Omicron variant and its subsequent subvariants. Nine mutations (5 in the heavy chain and 4 in the light chain) were introduced to yield pemivibart, which demonstrated greater breadth in neutralizing recent SARS-CoV-2 strains^3^. For example, data from the antibody manufacturer Invivyd, Inc. indicate that the engineered antibody could neutralize JN.1, the subvariant that dominated the first half of 2024, with 50% inhibitory concentrations (IC_50_) of 0.075 µg/mL and 0.063 µg/mL in pseudovirus and authentic virus neutralization assays in vitro, respectively^3^.

SARS-CoV-2 JN.1 subvariant first emerged in late 2023 and rapidly became dominant globally. In the past 6 months, JN.1 has continued to evolve, giving rise to multiple sublineages with unique spike mutations (**Figure S1A**). KP.2 was the first significant progeny to appear, but it was later outcompeted by KP.3, with both sublineages gradually displacing the original JN.1 (**Figure S1B**). More recently, KP.3.1.1, KP.2.3, and LB.1 have emerged, each independently developing a deletion of S31 (S31Δ) in the N-terminal domain (NTD) of spike (**Figure S1A**). Importantly, KP.3.1.1 is now the fasting growing sublineage worldwide (**Figure S1B**).

We therefore assessed the impact of recent SARS-CoV-2 evolution on the neutralizing activity of pemivibart. Pseudoviruses for JN.1, KP.2, KP.3, KP.2.3, KP.3.1.1, and LB.1 were constructed and subjected to neutralization assays as previously described^4^. Pemivibart neutralized both JN.1 and KP.2 in vitro with comparable activity, whereas its potency was decreased slightly against LB.1, KP.2.3, and KP.3 but substantially against KP.3.1.1 (**Figure 1A**). Critically, the IC_50_ of pemivibart against KP.3.1.1 was ∼6 µg/mL, or ∼32.7 fold higher than the IC_50_ for JN.1 in our direct comparison of the two viruses (**Figure 1B**). We note that our virus neutralization assays yielded results concordant to those reported by Invivyd, the manufacturer of pemivibart, against multiple SARS-CoV-2 strains (**Figure S2**).

**Figure 1.**
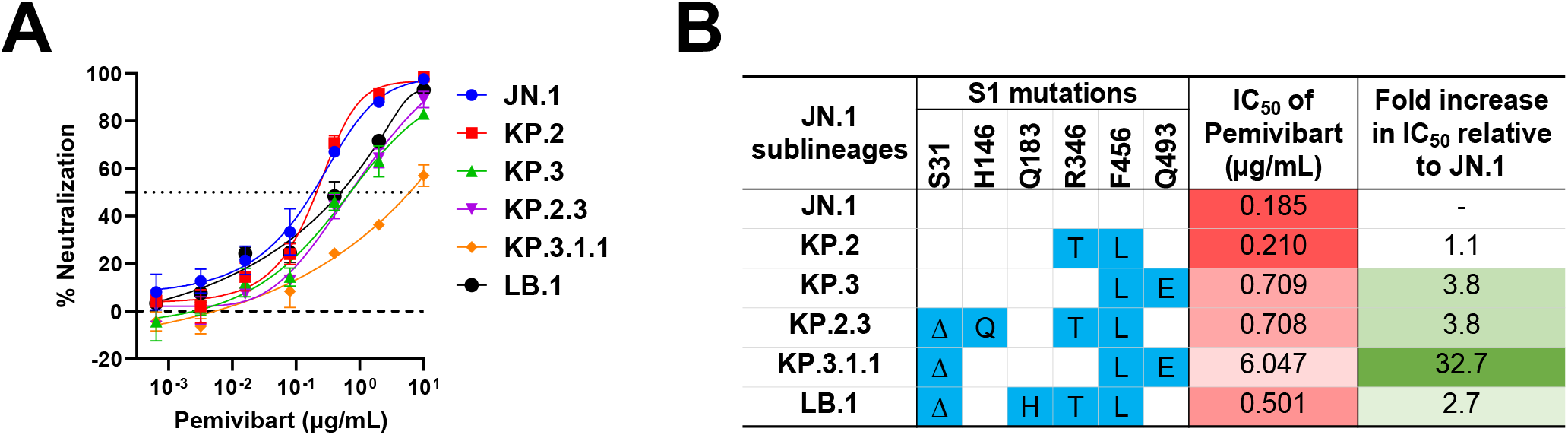
Neutralization of pemivibart against JN.1 sublineages. **A**. Neutralization curves of Pemivibart against the indicated pseudotyped viruses. **B**. Spike mutations in the S1 region, along with the corresponding neutralization IC_50_ values and fold-changes for the indicated JN.1 sublineages. Δ, deletion.

Comparing the results of KP.2 versus KP.3 (**Figure 1B**), it is apparent that Q493E mutation has a greater impact on pemivibart activity than R346T mutation. Although Q493E does not reside within the epitope of ADG-2^2^ (and presumably pemivibart), it may increase resistance to pemivibart due to an enhanced binding affinity for the ACE2 receptor facilitated by an epistatic effect with the F456L mutation^5^. Comparing the results of KP.3 versus KP.3.1.1 (**Figure 1B**), it is also apparent that S31Δ mutation has an even larger impact (∼9 fold) on the engineered antibody. Interestingly, multiple sublineages have converged to this mutation independently (**Figure S1A**). How S31Δ confers resistance to pemivibart, and presumably to serum antibodies in the population as well, is not immediately clear given it is quite distal to the typical SARS-CoV-2 neutralizing epitopes. We note, however, that this deletion results in the creation of a potential N-linked glycosylation site (**Figure S3A**), as well as the loss of hydrogen bonding with and F58, F59 and S60 (**Figure S3B**). Both changes could alter the conformation at the bottom of the NTD and thereby affect the up-and-down motion of the RBD (**Figure S3C**) that could negatively impact antibodies directed to so-called class 1 and class 1/4 regions of this domain. Data in **Figure S4** showed KP.3.1.1 with the S31Δ mutation was ∼8-fold more resistant to human ACE2 inhibition than KP.3, supporting the hypothesis that S31Δ may alter spike conformation and affect ACE2 engagement (**Figure S4**). The precise structural mechanism notwithstanding, our findings show a substantial loss of pemivibart activity against KP.3.1.1, the most rapidly expanding SARS-CoV-2 subvariant currently. Therefore, close monitoring of the clinical efficacy of this newly authorized monoclonal antibody is warranted.

## Supplementary Appendix

### Supplementary Methods

#### Cell lines

Vero-E6 (CRL-1586) cells and HEK293T (CRL-3216) cells were obtained from ATCC and cultured at 37°C with 5% CO_2_ in Dulbecco modified Eagle medium (DMEM) supplemented with 10% fetal bovine serum (FBS) and 1% penicillin-streptomycin. Expi293 cells (A14527) were purchased from Thermo Fisher Scientific and maintained in Expi293 expression medium per the manufacturer’s instructions. Morphology of each cell line was confirmed visually before use. All cell lines tested mycoplasma negative.

#### SARS-CoV-2 variant spike plasmids

Spike-expressing plasmids for JN.1, KP.2, and KP.3 were previously generated^1^. To make the spike-expressing plasmids for KP.2.3, KP.3.1.1, and LB.1 for pseudovirus production, spike mutations were introduced to the JN.1 spike-expressing construct using the QuikChange II XL site-directed mutagenesis kit (Agilent). Spike mutations of each subvariant on top of JN.1 are shown in **Figure S1A**.

#### Pseudovirus production

SARS-CoV-2 pseudoviruses were produced in a vesicular stomatitis virus (VSV) background, in which the native VSV glycoprotein was replaced by SARS-CoV-2 variant spikes, as previously described^2^. Briefly, the spike-expressing construct was transfected into HEK293T cells using 1mg/mL PEI MAX (Polysciences Inc). 24 hours later, the transfected cells were infected with VSV-G pseudotyped ΔG-luciferase (G*ΔG-luciferase, Kerafast) for 2 hours, then washed three times with culture medium before being cultured in fresh medium for an additional 24 hours. Anti-VSVG (I1) antibody^3^ was added to deplete non-pseudotyped viruses. Pseudoviruses were then harvested, clarified by centrifugation, aliquoted and stored at −80°C before use.

#### Antibody and hACE2 Expression and Purification

The Pemivibart antibody sequences for the heavy chain variable (VH) and light chain variable (VL) domains were obtained from the patent by Invivyd (WO2024050356A2), synthesized by GenScript, and subsequently cloned into the gWiz vector to generate antibody expression plasmids, as previously described^2^. To produce the Pemivibart antibody and human ACE2 (hACE2) with an Fc tag, plasmids encoding the heavy and light chains were co-transfected and the pcDNA3-sACE2-WT(732)-IgG1(Addgene #154104) plasmid was transfected into Expi293 cells using PEI-MAX. The cell supernatants were harvested after five days. The Pemivibart antibody and human ACE2 were then purified using Protein A Sepharose (Cytiva) according to the manufacturer’s instructions. Before use, the molecular weight and purity were confirmed by SDS-PAGE protein electrophoresis.

#### Pseudovirus neutralization and ACE2 inhibition assay

Each SARS-CoV-2 pseudovirus was titrated on Vero-E6 cells to standardize the viral infectious dose before use in neutralization assays or ACE2 inhibition assays. For neutralization assays, serially diluted antibodies (5-fold dilution starting from 10 µg/mL) were added to 96-well plates. Then, pseudoviruses were added and incubated with the serial antibody dilutions at 37 °C for 1 hour. Wells containing only pseudoviruses were included as controls in each plate. For ACE2 inhibition assays, hACE2 was diluted from 10 µg/mL with a dilution factor of three across seven serial dilutions. Following this, pseudoviruses were added and incubated with varying dilutions of either the serum or hACE2 at 37 °C for 1 hour. As a control, wells containing only the pseudovirus were also prepared on each test plate. Subsequently, Vero-E6 cells were then added at a density of 4 × 10^4^ cells per well and incubate at 37 °C overnight. Cells were lysed and luminescence was measured using the Luciferase Assay System (Promega) and SoftMax Pro v.7.0.2 (Molecular Devices) according to the manufacturers’ instructions. Data were analyzed using GraphPad Prism v.9.3.

## Acknowledgements

This study was supported by funding from the NIH SARS-CoV-2 Assessment of Viral Evolution (SAVE) Program (subcontract no. 0258-A709-4609 under federal contract no. 75N93021C00014) and the Gates Foundation (project INV019355) to D.D.H. and internal startup funding UR014016 from Columbia University to Y.G. We thank all who contributed their data to GISAID.

## Author Contributions

D.D.H. conceived of this project. Q.W. and J.H. performed experiments. Y.G. conducted bioinformatic analyses. Q.W., Y.G., and D.D.H. analyzed the results and wrote the manuscript.

## Supplementary Figures

**Figure S1.**
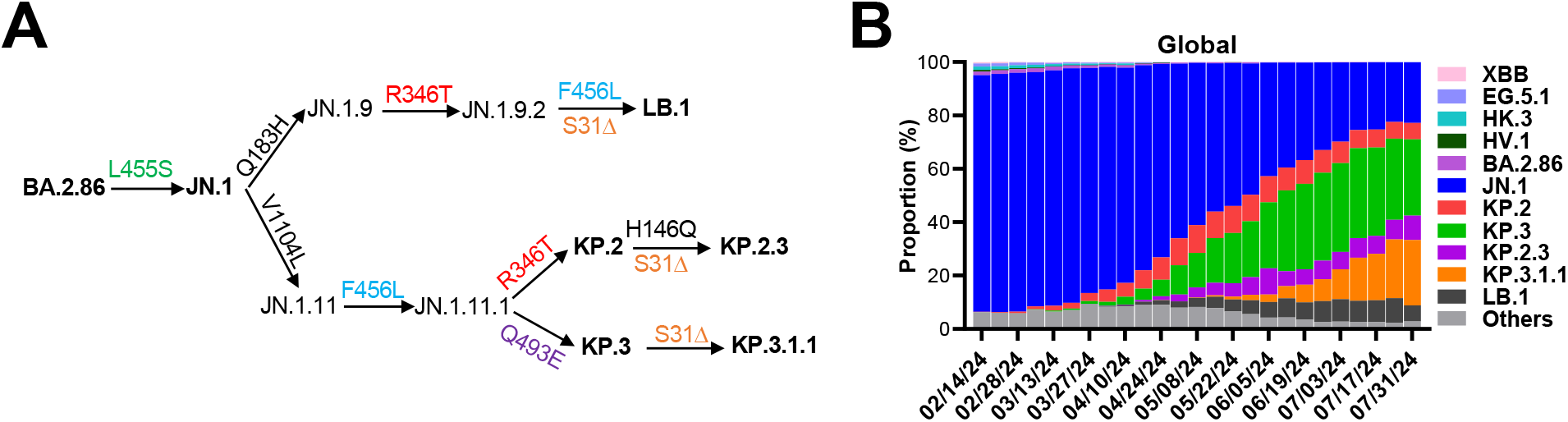
Overview of circulating SARS-CoV-2 JN.1 sublineages. **A**. Evolutionary pathway of JN.1 sublineages. Δ, deletion. **B**. Global frequencies of major SARS-CoV-2 forms during the indicated time period. Sequence frequencies were obtained from the Global Initiative on Sharing All Influenza Data (GISAID).

**Figure S2.**
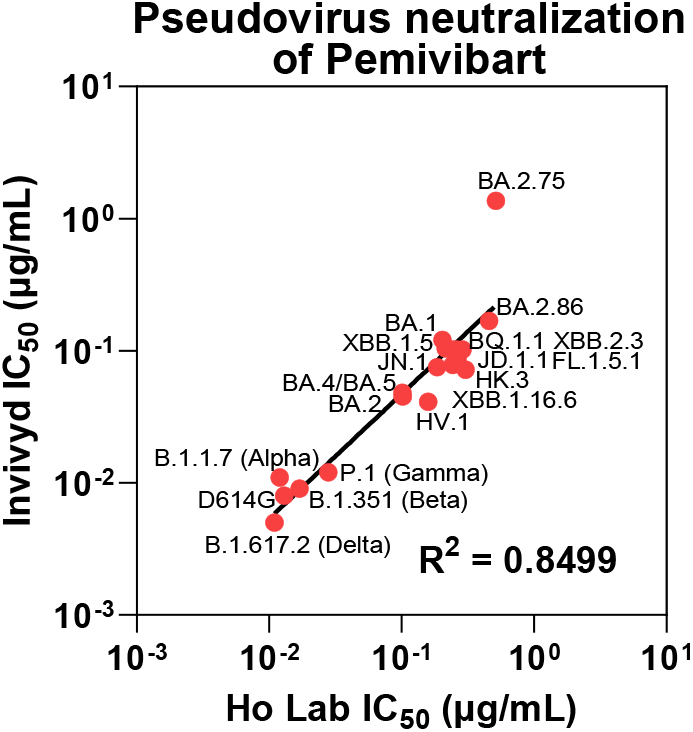
Correlation of pseudovirus neutralization IC_50_ values of pemivibart reported by Invivyd and values generated as a part of this study.

**Figure S3.**
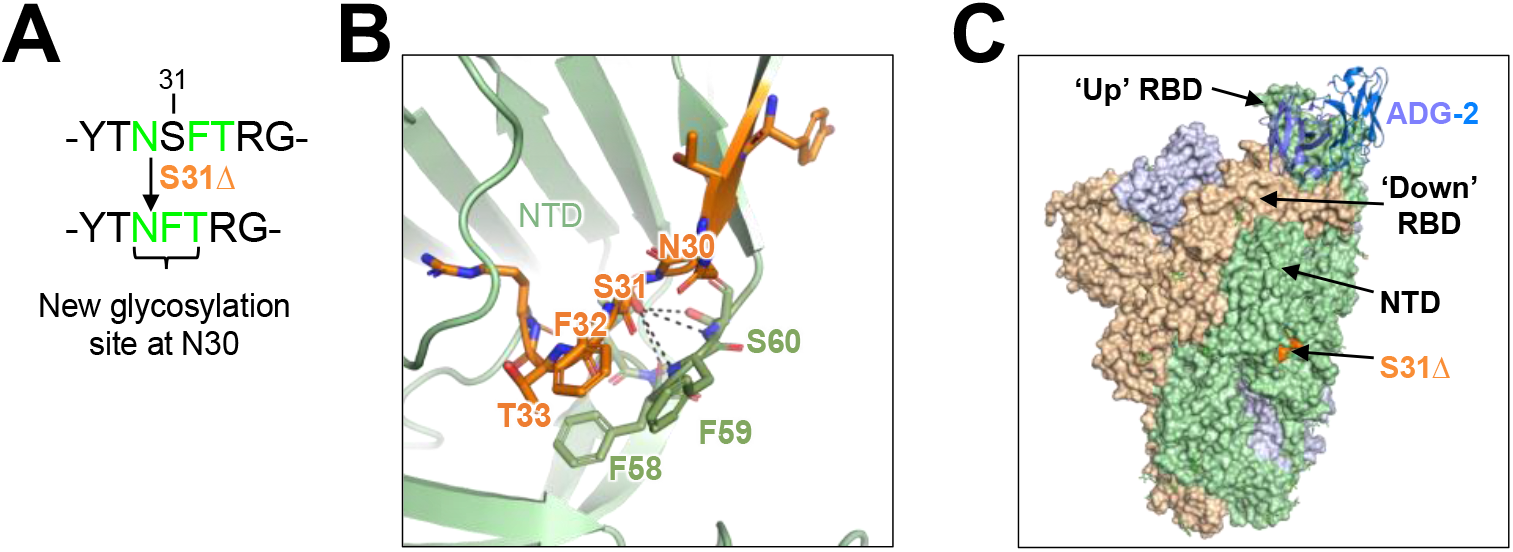
Predicted effects of the S31Δ mutation on the JN.1 spike. **A**. The S31Δ mutation may introduce a glycosylation site at N30. **B**. Structural details of the S31-surrounding region in the NTD. Black dash lines indicate hydrogen bonds. PDB: 8Y5J. **C**. Overall view of ADG-2/pemivibart binding site on the RBD in the up position and S31Δ on the external surface of NTD.

**Figure S4.**
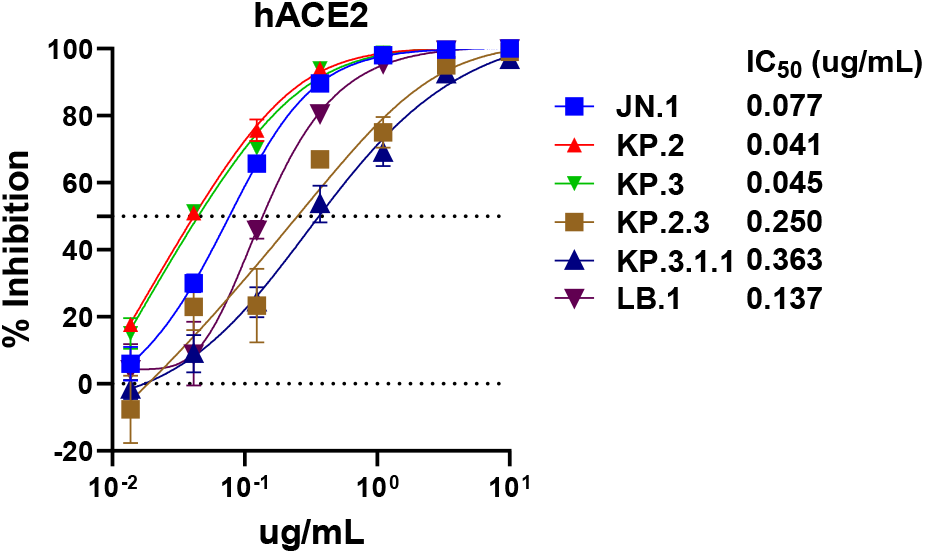
Susceptibility of JN.1 sublineage pseudoviruses to human ACE2(hACE2) inhibition on Vero-E6 cell with IC_50_ values denoted.

## References

1. FDA Roundup: March 22, 2024. 2024. (Accessed 07/25/2024, at https://www.fda.gov/news-events/press-announcements/fda-roundup-march-22-2024.)

2. Rappazzo CG, Tse LV, Kaku CI, et al. Broad and potent activity against SARS-like viruses by an engineered human monoclonal antibody. Science 2021;371:823–9.

3. CDER Scientific Review Documents Supporting Emergency Use Authorizations for Drug and Biological Therapeutic Products | COVID-19. 2024. (Accessed 08/10/2024, at https://www.fda.gov/drugs/coronavirus-covid-19-drugs/cder-scientific-review-documents-supporting-emergency-use-authorizations-drug-and-biological.)

4. Wang Q, Mellis IA, Bowen A, et al. Recurrent SARS-CoV-2 spike mutations confer growth advantages to select JN.1 sublineages. bioRxiv 2024:2024.05.29.596362.

5. Taylor AL, Starr TN. Deep mutational scanning of SARS-CoV-2 Omicron BA.2.86 and epistatic emergence of the KP.3 variant. bioRxiv 2024.

## Supplementary References

1. Wang Q, Mellis IA, Bowen A, et al. Recurrent SARS-CoV-2 spike mutations confer growth advantages to select JN.1 sublineages. bioRxiv 2024:2024.05.29.596362.

2. Liu L, Wang P, Nair MS, et al. Potent neutralizing antibodies against multiple epitopes on SARS-CoV-2 spike. Nature 2020;584:450–6.

3. Lefrancios L, Lyles DS. The interactionof antiody with the major surface glycoprotein of vesicular stomatitis virus. I. Analysis of neutralizing epitopes with monoclonal antibodies. Virology 1982;121:157–67.

